# Light Microscopy-Based Organelle Quantification: A Comprehensive Protocol

**DOI:** 10.64898/2026.01.19.700276

**Authors:** Suraj Thapilyal, Namdar Hari Kalpana, McMillan Ronald, Jermiah Afolabi, Andrea Marshall, Prasanna Venkhatesh, Ravi Kumar Pujala, Antentor O Hinton, Hailey Parry, Brian Glancy, Prasanna Katti

## Abstract

Cellular organelles are not just static structures; they are highly dynamic and directly linked to cellular functions. Changes in their morphology can be early indicators of diseases. Recent advancements in light microscopy techniques have transformed organelle research from qualitative descriptions to precise, quantitative measurements, enabling nanoscale resolution, high-throughput image analysis, and live-cell compatibility. This enables accurate measurement of organelle morphology, dynamics, and spatial organization using modern imaging and analysis techniques. By quantifying organelles, we go beyond simply visualizing to measuring and statistically comparing cellular features across different samples. This protocol addresses a wide range of cellular organelles across all major experimental systems, specifically mentioning mitochondria, myofibers, actin filaments, endoplasmic reticulum, and Golgi apparatus, by integrating experimental design, optimized sample preparation, high-resolution imaging, and validated Fiji/ImageJ-based analysis workflows. For each organelle, step-by-step methods specify reagents, equipment, acquisition parameters, and expected results. While recent advances, such as expansion microscopy, correlative light-electron microscopy, and AI-powered segmentation, offer gains in throughput and resolution, this workflow demonstrates that Fiji-based analysis remains fully capable of delivering high-precision organelle quantification. The entire workflow can be completed within 2-4 weeks, from initial design through validation and the production of measurements suitable for cross-study comparisons. Overall, this protocol establishes a flexible approach to standardize organelle quantification to understand multiple organelles simultaneously in their cellular contexts.

Basic Protocol 1: Mitochondrial Quantification

Basic Protocol 2: Myofibril Quantification

Basic Protocol 3: Golgi Apparatus Morphometry

Basic Protocol 4: Endoplasmic Reticulum Network Analysis

Alternate Protocol 1: Super-Resolution Imaging Protocol

## Introduction

Cellular organelles are discrete, functionally specialized compartments that orchestrate the complex biochemical processes essential to life (Glancy et al. 2020; Zulueta Diaz and Arnspang 2024). Quantitative analysis of organelle morphology, dynamics, and spatial organization has become increasingly critical for understanding cellular physiology, disease mechanisms, and therapeutic interventions. The past decade has witnessed revolutionary advances in light microscopy techniques, enabling researchers to achieve nanoscale resolution while maintaining compatibility with live-cell imaging and high-throughput analysis (Bond et al. 2022; Katti et al. 2022; Schmied et al. 2024). Super-resolution microscopy achieves sub-100 nm resolution (Vicidomini et al. 2018; Zulueta Diaz and Arnspang 2024). Artificial intelligence-powered image analysis with human-level accuracy (Heinrich 2021; Morone et al. 2020). Correlative approaches bridging light and electron microscopy (Heinrich 2021) advanced labelling strategies enabling multiplexed visualization, and automated analysis pipelines supporting large-scale studies. These advances have transformed organelle research from qualitative descriptions to precise, quantitative measurements, revealing fundamental cellular organization principles that are standardized and established in electron microscopy and evident in the universal TEM analysis workflow. This workflow provides essential guidelines for light microscopy-based analysis (Lam et al., 2021). This protocol addresses the full spectrum of cellular organelles across all major experimental systems, including mitochondria, myofibers, Actin filaments, endoplasmic reticulum, and Golgi apparatus. The methodologies described here incorporate recent cutting-edge developments, including deep learning segmentation tools, microscopy protocols achieving 30 nm resolution, and analysis pipelines that process datasets within hours rather than weeks (Morone et al. 2020).

## Strategic Planning Considerations

### Experimental Design Framework

Formulating biological questions represents the critical first step in organelle quantification studies. Different research objectives require distinct methodological approaches: morphological studies benefit from high-resolution structural imaging, functional analyses require multi-parameter measurements combining structural and biochemical readouts, and dynamic studies demand high temporal resolution with minimal photodamage. Sample selection must account for cell-type-specific organelle characteristics. Cultured cell lines offer standardized conditions and genetic manipulability but may not recapitulate physiological organelle organization. Primary cells maintain native characteristics but introduce experimental variability (Al-Kofahi et al., 2018). Tissue sections preserve spatial relationships but require specialized processing protocols.

### Technology Selection Matrix

Imaging modality selection depends on resolution requirements, sample compatibility, and throughput needs. Confocal microscopy provides excellent 3D imaging capabilities for routine morphological analysis. Super-resolution techniques, such as STED microscopy, achieve resolutions of 50-90 nm, which is essential for detailed structural studies (Bond et al., 2022; Prakash et al., 2025). Expansion microscopy democratizes super-resolution capabilities by achieving effective resolutions of 60-70 nm on standard confocal systems (Gao et al., 2018). Analysis complexity scales with biological questions. Simple counting applications can utilize traditional thresholding approaches, while morphological analysis requires advanced segmentation algorithms. Dynamic studies require particle-tracking capabilities, and multi-organelle interactions necessitate sophisticated colocalization analysis.

### Validation Strategy Framework

Multi-modal validation ensures biological relevance and technical accuracy. Light microscopy quantification should be validated against electron microscopy gold standards, where possible (McCafferty et al., 2024). Functional assays must correlate with morphological measurements. Prior EM studies have shown the effects of inconsistent measurement practices. Standardized sampling strategies are therefore essential. Clear segmentation and validation are crucial for light microscopy-based organelle morphometry (Neikirk, Lopez, et al., 2023a). Pharmacological perturbations provide positive and negative controls for method validation.

## Basic Protocols for Major Organelle Types

### Protocol 1: Mitochondrial Quantification

**Objective**: Comprehensive morphological and functional analysis of mitochondrial networks in *Drosophila* muscle tissue, followed by Fiji-based quantification of morphology and fragmentation (Figure 1).

**Figure 1.**
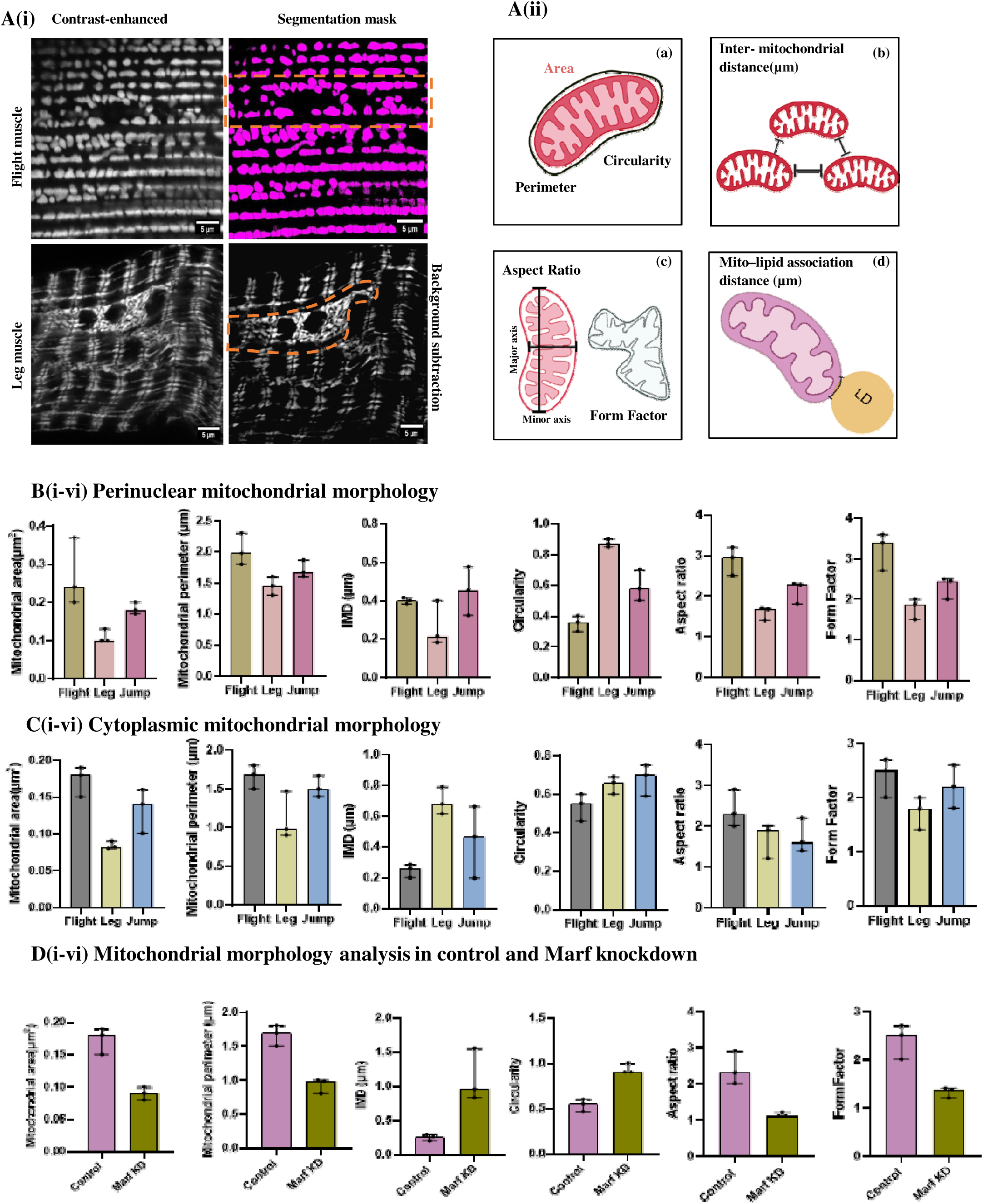
Quantitative analysis of Mitochondrial morphology in Drosophila Flight, Leg and Jump muscles through Fiji. (A(i)) Representative images showing *Drosophila* flight and leg muscle segmented mitochondria: Contrast-enhanced image (left), and Fiji segmentation overlay (top right) for flight muscle, with background subtracted for leg muscle (bottom right),Dotted regions represent perinuclear mitochondria. (A(ii)) Schematic illustrating mitochondrial morphology parameters, including (a) area, perimeter, and circularity; (b) inter-mitochondrial distance (IMD); (c) aspect ratio and form factor; and (d) lipid–mitochondria association distance. (B) Quantitative analysis of perinuclear mitochondria in flight and leg muscles. (C) Quantitative analysis of cytoplasmic mitochondria in flight and leg muscles. (D) Representative examples of mitochondrial morphology in control and Marf knockdown conditions. Statistical comparisons among three groups were performed using the Kruskal–Wallis test and for two groups the Mann–Whitney U test. Data are presented as medians with interquartile ranges. For B–D (i–vi), each dot represents one independent image (n = 3 images per condition). The number of mitochondria quantified per image was as follows: C (perinuclear)-Flight: 61, 15, 51; leg: 62, 90, 67; jump: 56, 19, 15; D (cytoplasmic)-Flight: 206, 197, 165; leg: 185, 147, 176; jump: 136, 190, 78; E- Control: 223, 196, 165; Marf KD: 275, 268, 198 mitochondria. (E) Representative mitochondria–lipid droplet images are shown alongside quantification of inter-mitochondrial distance, mitochondrial area, perimeter, aspect ratio, and circularity. Data are presented as median with interquartile range. Statistical comparisons between two groups (n = 101 and n = 43) were performed using the Mann–Whitney U test, with individual data points representing independent measurements. Figure.1E(ii), n=43, n represents no. of mitochondria lipid droplet associations

**Materials**

**Reagents**

1. Mito-GFP line (BDSC-95270), Mef2-gal4(BDSC-67041)
2. MitoTracker Red CMX Ros (Invitrogen, M7512)
3. Phosphate-buffered saline (PBS), pH 7.4
4. PBST (Triton X-100)
5. 4% Paraformaldehyde (PFA)
6. TMRM (Tetramethyl rhodamine methyl ester)
7. Mounting medium: ProLong Gold or similar anti-fade reagent

**Equipment**

Confocal microscope with 63x/1.4 NA oil objective minimum

Incubation chamber with CO and temperature control

**Procedure**

**Sample preparation** (45 minutes):

1. Culture cells on glass coverslips to 60-80% confluence.
2. For live imaging: Load with 250 nM MitoTracker Red for 30 minutes at 37°C.
3. For fixed samples: Fix with 4% paraformaldehyde for 10 minutes, permeabilize with 0.1% Triton X-100.
4. For Muscle samples: Fix with 4% paraformaldehyde for 1-1.5 hr, permeabilize with 0.1% Triton X-100.

**Image acquisition**

5 Acquire z-stacks with 0.1-0.2 μm spacing.
6 Use sequential scanning to minimize bleed-through.
7 Maintain consistent exposure settings across samples.
8 Image minimum 30 cells per condition across multiple fields.

**Quantitative analysis**

9 Preprocess images with deconvolution if available.
10 Extract morphological parameters: volume, length, connectivity.
11 Calculate fragmentation indices and network metrics.

**Expected Results**

Morphological parameters typically range from 15-40% mitochondrial volume fraction in cultured cells/ Drosophila muscle tissue. Network connectivity shows PHI ratios of 0.6-0.9 in healthy cells, decreasing to 0.1-0.3 upon fragmentation-inducing treatments. Validation controls treated with FCCP should show significant fragmentation within 1-2 hours.

### Protocol 2: Lipid Droplet Identification and Image Processing

**Objective:** Identification and quantitative isolation of lipid droplets in isolated mouse skeletal muscle fibers using confocal microscopy and machine learning–assisted image analysis.

**Materials**

Animals-C57BL/6 mice

**Reagents**

1. Collagenase D (3 mg/mL)
2. Lipi Deep Red (200 nM)
3. Appropriate physiological buffer (e.g., Tyrode’s or PBS)
4. Software
5. ImageJ/Fiji
6. ilastik (pixel classification workflow)

**Procedure**

**Sample Preparation and Cell Isolation**

1. Dissect FDB muscle from C57BL/6 mouse hindlimb
2. Digest in 3 mg/mL collagenase D at 37 °C for 1 hour
3. Replace enzyme with fresh buffer
4. Isolate single muscle fibers by gentle trituration

**Lipid Droplet Staining**

1. Stain isolated fibers with 200 nM Lipi Deep Red for 30 min (RT)
2. Wash with fresh buffer

**Image Acquisition**

1. Acquire images on Zeiss LSM 980 confocal microscope
2. Use 63x oil-immersion objective
3. Maintain identical imaging settings across samples

**Image Processing and Lipid Droplet Identification**

1. Save Lipi Deep Red channel as TIF in ImageJ/Fiji
2. Perform pixel classification in plastic to identify lipid droplets
3. Export prediction map and generate a binary mask
4. Multiply mask with original image to isolate lipid droplet signal

**Quantitative Analysis**

1. Lipid droplet number
2. Area / volume
3. Spatial distribution within muscle fibers

### Protocol 3: Myofibril Quantification

**Objective**: Morphological analysis of myofibril organization in *Drosophila* muscle tissue, followed by Fiji-based quantification.

**Materials**

**Reagents**

Phalloidin- FITC/ TRITC

**Procedure**

**Sample preparation** (45 minutes):

1. Dissect *Drosophila* muscle tissue (IFMs) in cold PBS (Rai M et al., 2014).
2. Sample fixation with 4% PFA for 1.5 hr at room temperature.
3. Wash sample 3 times with PBS, 15 min each, Permeabilize with 0.1% Triton X-100.

**Phalloidin staining (direct F-actin labeling):**

4 Incubate samples with Phalloidin for 30 mins at RT in the dark.
5 Wash samples 3 times with PBS, 15 min each.
6 Mount using ProLong Gold or a similar anti-fade reagent.

**Image acquisition**

7 Acquire z-stacks with 0.2 μm spacing.
8 Use sequential scanning to minimize bleed-through.
9 Maintain consistent exposure settings across samples.
10 Image minimum 30 cells per condition across multiple fields.

**Quantitative analysis**

11 Preprocess images with deconvolution.
12 Fiji/ ImageJ analysis- Measure sarcomere length and width by identifying striation patterns in phalloidin-stained myofibrils through MyofibrilJ.

**Expected Results**

Average sarcomere lengths typically range from 2.5 µm to 3.77 µm in healthy *Drosophila* muscle fibers.

### Protocol 4: Golgi Apparatus Morphometry

**Objective**: Quantitative analysis of Golgi ribbon structure and fragmentation states.

**Materials**

**Reagents**

Transgenic Fly lines (RFP targeting Golgi Complex)

Optional: BODIPY-ceramide for metabolic labeling

**Procedure**

**Multi-marker staining**

1. Sequential staining protocol to minimize cross-reactivity.

**3D/2D acquisition**

2 Z-spacing: 0.1-0.15 μm for detailed ribbon morphology.
3 Super-resolution (3D-SIM or Airyscan) is preferred for fine structure.

**Morphological quantification**

4 Automated segmentation with morphological filters.
5 Extract volume, surface area, and fragmentation indices.
6 Analyze ribbon continuity and perinuclear distribution.

**Expected Results**

Healthy Golgi exhibits compact, perinuclear ribbons with 2-8 μm³ volumes. Fragmentation treatments increase the number of discrete Golgi elements while reducing average size.

### Protocol 5: Endoplasmic Reticulum Network Analysis

**Objective**: Three-dimensional reconstruction and quantification of ER tubular networks and cisternal structures.

**Materials**

**Reagents**

ER-Tracker Red (Invitrogen, E34250) or UAS-KDEL-RFP (**BDSC- 30909 and 30910**)

Primary antibodies: Calnexin, KDEL

**Procedure**

**Network labelling**

1. For live imaging: ER-Tracker Red (1 μM) for 30 minutes.
2. For immunofluorescence: Calnexin primary antibody (1:500).
3. For muscle tissues (used transgenic-KDEL tagged RFP).

**High-resolution imaging**

4 Use a 60x/1.4 NA objective for optimal network resolution.
5 Z-spacing: 0.1 μm for complete 3D network capture.

**Network analysis**

6 Employ the AnalyzER software for automated tubule extraction.
7 3D reconstruction and segmentation (Neikirk, Vue, et al., 2023).
8 Quantify branch points, total length, and cisternal areas.
9 Calculate network connectivity indices.

**Expected Results**

ER networks typically exhibit 200-500 branch points per cell with total tubule lengths of 50-200 μm. Cisternal regions comprise 20-40% of total ER volume in most cell types.

## Alternate Protocols for Specialized Applications

### Super-Resolution Imaging Protocol

**Applications**: Sub-diffraction analysis of organelle ultrastructure and protein localization.

**Critical Parameters**

Laser optimization, balance depletion efficiency with photobleaching. Time-gated detection with 0.5-6 ns windows improves resolution and reduces background.

**Procedure Modifications**

1. Fluorophore selection: ATTO series, Alexa Fluor series (Vicidomini et al., 2018).
2. Sample preparation: Enhanced fixation protocols for better structural preservation.
3. Acquisition optimization: Pixel size 20-30 nm for adequate sampling.
4. Deconvolution processing: Apply Richardson-Lucy or Wiener filtering.

### Equipment and Software Requirements

#### Essential Microscopy Systems

Confocal microscopy represents the minimum requirement for 3D organelle quantification. Modern systems should include multiple laser lines (405, 488, 561, 633 nm), high-sensitivity detectors (GaAsP preferred), and precision z-control (±10 nm accuracy) (Elliott, 2019). Super-resolution capabilities expand analytical possibilities. STED systems provide 50-90 nm resolution with live-cell compatibility. Structured illumination microscopy offers a 2X improvement in resolution with full-field imaging. Single-molecule localization techniques achieve 10-20 nm precision for specialized applications (Kumar et al., 2024; Prakash et al., 2025).

Environmental control proves essential for live-cell studies. Temperature stability (±0.1°C), CO regulation (±0.1%), and humidity control maintain physiological conditions during extended imaging sessions.

#### Software Analysis Platforms

Open-source solutions provide powerful, customizable analysis capabilities. ImageJ/FIJI, with specialized plugins, offers comprehensive organelle analysis tools (Neal et al., 2024; Schindelin et al., 2012). CellProfiler enables high-throughput quantification with automated workflows. Python-based frameworks such as CellPose and StarDist incorporate state-of-the-art deep learning methods. Commercial platforms deliver integrated analysis environments. IMARIS provides advanced 3D visualization and quantification. Columbus supports high-content screening applications. Huygens offers professional deconvolution and super-resolution processing. AI-powered tools represent the cutting edge of automated analysis. LysoQuant achieves human-level accuracy for endolysosome segmentation. MitoGraph provides specialized mitochondrial network analysis (Viana et al., 2015). Deep learning approaches increasingly outperform traditional segmentation methods (Fischer et al., 2020; Morone et al., 2020).

## Step-by-Step Procedures

### Standard Workflow for Organelle Quantification

**Phase 1: Experimental Planning (1-2 weeks)**

Define a biological question and select appropriate organelle targets.

Choose an imaging modality based on resolution and throughput requirements. Validate antibodies and labelling protocols for target organelles.

Establish an analysis pipeline and validation criteria.

Calculate sample sizes using power analysis for statistical significance.

**Phase 2: Sample Preparation (1-3 days)**

**Cell culture preparation**

1. Culture cells on glass coverslips or imaging dishes.
2. Maintain consistent culture conditions across all samples.
3. Time experimental treatments to coincide with optimal cell density.

**Labeling protocols**

1. For live imaging: Apply organelle-specific trackers 30-60 minutes before imaging.
2. For fixed samples: Optimize fixation conditions (4% PFA, 10-15 minutes typical).
3. Perform immunofluorescence using validated fluorophores/antibody concentrations.

**Quality control**

1. Assess labelling specificity using appropriate controls.
2. Verify cell viability for live imaging applications.
3. Document any morphological abnormalities.

**Phase 3: Image Acquisition (1-5 days)**

**System calibration**

1. Perform daily laser alignment and detector optimization.
2. Verify focus stability using reference samples.
3. Calibrate spatial measurements using fluorescent beads.

**Acquisition parameters**

4 Set consistent exposure times and gain settings.
5 Use appropriate z-spacing for 3D reconstruction (typically 0.1-0.2 μm).
6 Image sufficient cells for statistical analysis (minimum 30 per condition).

**Data management**

7 Implement consistent file naming conventions.
8 Backup raw data immediately after acquisition.
9 Document acquisition parameters for reproducibility.

**Phase 4: Image Analysis (2-7 days)**

**Preprocessing**

1. Apply noise reduction and deconvolution as appropriate.
2. Perform intensity normalization across samples.
3. Generate maximum intensity projections for 2D analysis if needed.

**Segmentation and quantification**

4 Apply validated segmentation algorithms.
5 Extract morphological and functional parameters.
6 Implement quality control measures for automated analysis.

**Data validation**

7 Compare automated results with manual analysis on a subset of data.
8 Apply appropriate statistical tests for biological questions.
9 Generate summary statistics and visualizations.

### Critical Parameters and Troubleshooting

#### Common Technical Issues

Poor image quality is the most common obstacle to successful quantification. Sample preparation significantly affects the observed shape of organelles; fixation strength, permeabilization, and labeling conditions can create structural artifacts (Hinton Jr. et al., 2023). Possible solutions include optimizing sample preparation protocols, adjusting imaging parameters to improve signal-to-noise ratios, and implementing appropriate image preprocessing algorithms. Segmentation errors can significantly impact quantitative results. Mitigation strategies consist of validating segmentation against manual annotations, adjusting threshold parameters for specific samples, and employing multiple segmentation techniques for cross-validation. Photobleaching and phototoxicity restrict live-cell imaging applications. Recommended approaches are minimizing laser power while ensuring sufficient signal, using appropriate anti-fade reagents, and applying time-gated acquisition strategies.

#### Validation Strategies

Biological validation ensures that measured parameters accurately reflect the true properties of organelles. Use established pharmacological treatments as positive controls (e.g., FCCP for mitochondrial fragmentation and brefeldin A for Golgi disruption). Implement genetic perturbations with known phenotypes for method validation. Technical validation confirms the accuracy and precision of measurements. Compare results across different imaging modalities and analysis software. Conduct inter-operator reproducibility studies. Validate against electron microscopy whenever possible.

#### Quality Control Metrics

Signal-to-noise ratios should exceed 3:1 for quantitative analysis. Resolution measurements using decorrelation analysis ensure adequate sampling. Photon-count statistics verify the presence of sufficient signals for reliable measurements.

### Statistical Analysis and Data Presentation

#### Experimental Design Recommendations

Sample size calculations require careful consideration of expected effect sizes and measurement variability. Power analysis should target 80% power to detect biologically meaningful differences. Nested data structures (multiple organelles per cell, multiple cells per condition) necessitate suitable statistical models. Control selection is crucial for interpretable results. Include negative controls (unstained samples, secondary antibody only) and positive controls (known organelle modulators). Blinding procedures reduce analysis bias, which is especially important for manual measurements.

#### Statistical Methods

Multiple comparison corrections prevent false discoveries when testing multiple hypotheses. Apply Bonferroni correction for small numbers of comparisons or false discovery rate control for larger testing scenarios. Effect size reporting provides a biological context beyond statistical significance. Cohen’s d or similar measures quantify the magnitude of observed differences, allowing for the assessment of biological relevance.

#### Data Visualization Best Practices

Representative image selection should follow unbiased criteria rather than highlighting extreme examples. Include scale bars, proper contrast adjustment, and consistent presentation parameters across conditions—quantitative data presentation benefits from showing distributions rather than summary statistics alone. Box plots reveal the spread of data and outliers. Violin plots combine density information with traditional box plot elements. Statistical graphics should indicate sample sizes, the meanings of error bars (standard error vs. standard deviation), and the results of statistical tests. Report confidence intervals alongside point estimates where appropriate.

## Understanding Figures

**Figure 1. Quantitative analysis of Mitochondrial morphology in *Drosophila* Flight, Leg, and Jump muscles through Fiji**

Mitochondrial structural parameters were analyzed in *Drosophila* flight and leg muscles. As observed in (Figure 1A), mitochondria in flight muscles are compactly arranged along myofibrils, whereas mitochondria are more dispersed with a less ordered arrangement in leg muscle. All quantitative analyses were performed using Fiji-based image analysis to ensure reliable measurement and objectivity.

Mitochondria were categorized as perinuclear and cytoplasmic based on their location relative to the nucleus. Through a segmentation-based analysis pipeline, several important parameters, including area, perimeter, Inter-mitochondria distance, aspect ratio, circularity, and form factor, were extracted. Mitochondrial area and perimeter were calculated to describe the size and boundary length of each mitochondrion. These measurements provide insight into the overall mitochondrial dimensions within the observed area (Figure 1B and 1C). Inter-mitochondral distance (IMD) was calculated as the distance between the nearest mitochondrial centroids. Circularity assessed how closely mitochondrial profiles resembled circle by considering both area and perimeter. A circularity value of 1 indicates a perfect circle. Lower values correspond to stretched or irregular shapes. This measure is commonly used to evaluate changes in mitochondrial shape and fragmentation. To describe mitochondrial shape, the aspect ratio quantified degree of elongation along the major axis. Higher aspect ratios indicate elongated structures, while values closer to 1 indicate rounder shapes. Form factor indicated boundary irregularity and complexity, reflecting deviations from smooth, round profiles. Lower form factor values indicate more irregular boundaries, often linked to fragmented or highly branched structures. Inter-mitochondral distance (IMD) was calculated as the distance between the nearest mitochondrial centroids.

Marf (Mitochondrial assembly regulatory factor) is the *Drosophila* counterpart of Mitofusin. It plays an important role in the fusion of the outer mitochondrial membrane. This process is essential for maintaining a connected network of mitochondria and their normal shape (Katti P et al., 2021, Katti P et al., 2022). Images from experiments in which Marf was knocked down showed changes in mitochondrial shape compared with controls. The mitochondria appeared more rounded and fragmented. These examples highlight how disrupting mitochondrial fusion can lead to noticeable changes in morphometric parameters (Figure 1D). Though Marf knockdown was not the main focus of this study, it provides a biological reference. This helps confirm that the morphological measurements used are sensitive and relevant for detecting changes in mitochondrial structure.

Figure 1E(i) shows an image of mitochondria and lipid droplets, along with a quantitative assessment of mitochondrial arrangement and shape. We measured the distance between mitochondria and lipid droplet contact. We assessed the morphological parameters of mitochondria, which are associated with lipids and non-associated with lipids Figure 1E(ii-vi). Together these findings provide a consistent method to evaluate mitochondrial shape and arrangement in relation to lipid droplets.

These metrics provide a clear framework for evaluating mitochondrial size, complexity, abundance, and spatial arrangement and demonstrate that mitochondrial networks are specialized within different muscle types.

Figure 2: Lipid Droplet Organization and Morphology in Muscle Fibers

Figure 2 shows how to identify, segment, and quantitatively describe lipid droplets in muscle fibers. Panel A displays a confocal image of lipid droplets scattered within muscle fibers. This highlights their punctate organization along the structure of the myofibrils. The corresponding segmented image shows clear detection of individual lipid droplets using the image processing pipeline. Each droplet is distinctly separated from the background. Panels B(i–iii) illustrate the distribution of lipid droplet shape parameters, including area, perimeter, and circularity. These graphs show the variety in lipid droplet size and form, reflecting differences in lipid storage characteristics at the single-droplet level. Overall, this figure demonstrates the reliability of the segmentation method and the ability to quantitatively describe lipid droplet shape in muscle fibers using light microscopy analysis.

**Figure 2.**
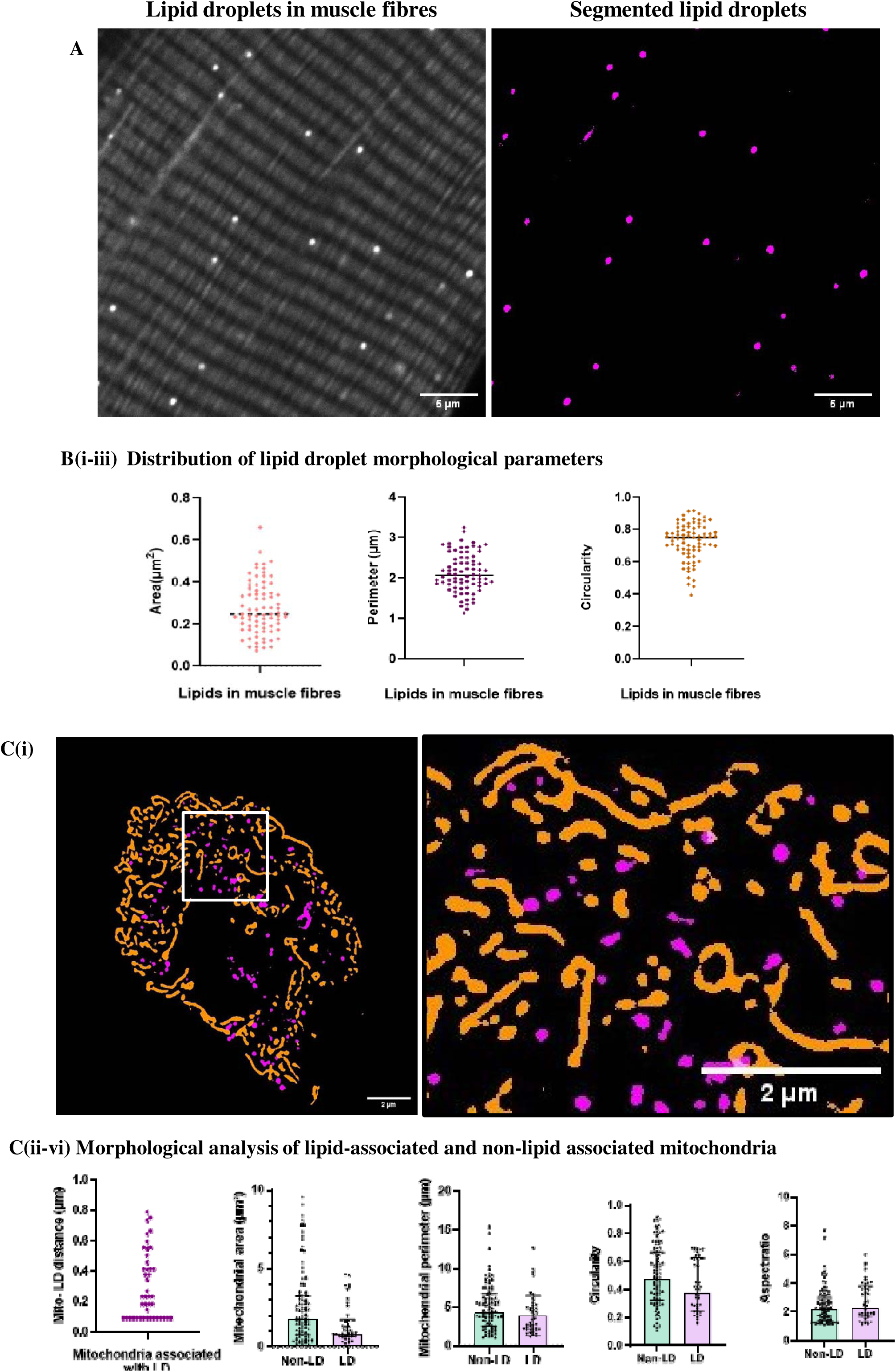
Quantification of lipid droplets in muscle fibres. (A)Representative image of lipid droplets in muscle fibres and corresponding segmented mask are shown.(B) Scatter plots depict lipid droplet area, perimeter, and circularity quantified from n=80 lipid droplets pooled from 3 independent images, with individual data points representing independent lipid droplets and central lines indicates median. (C) Representative mitochondria–lipid droplet image is shown(i), alongside quantification of mitochondrial- lipid association distance, area, perimeter, aspect ratio, and circularity (ii-vi) respectively. Data are presented as median with interquartile range. Statistical comparisons between two groups (n = 101 and n = 43) were performed using the Mann–Whitney U test, with individual data points representing independent measurements. Figure.2C(ii), n=43, n represents no. of mitochondria lipid droplet associations

**Figure 3. Morphometric measurements of *Drosophila* Flight muscle sarcomeres using MyofibrilJ.**

The contractile machinery of the *Drosophila* flight muscle consists of uniform, cylindrical myofibrils extending along the entire length of the muscle fibers. Each myofibril is composed of hundreds of serially connected, nearly uniform-sized sarcomeres (Figure 3A). Consequently, sarcomere length and myofibril width (or diameter) are frequently reported morphometric traits. Despite the remarkable regularity of sarcomeres, the average sarcomere lengths range from 2.9µm to 3.77 µm in adult flies (Görög et al., 2025). To determine these parameters, we used the MyofibrilJ plugin in Fiji, which relies on Fast Fourier Transform (FFT)- based analysis to identify repeating striation patterns.

**Figure 3.**
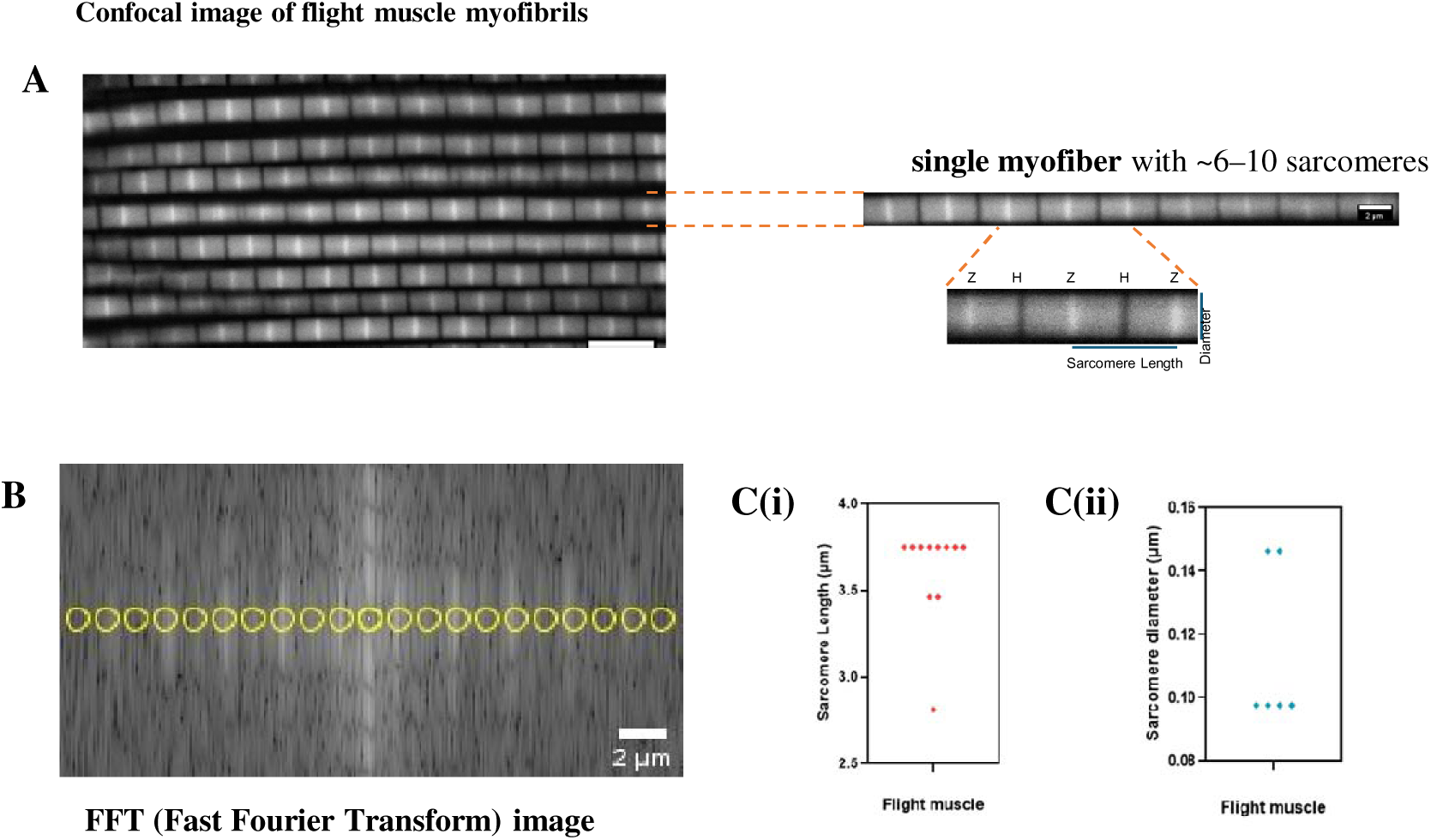
Morphometric measurements of Drosophila Flight muscle sarcomeres using MyofibrilJ. (A)Confocal micrograph showing flight muscle myofibrils with sarcomere arrangement. Zoomed view of a single myofibril consists of regular, uniform sarcomeres with characteristic lengths and diameters. The Z-discs (Z) at the borders of sarcomeres and the central H-zone (H) are labelled. Sarcomere length is determined from images where the Z-disc is labelled, while diameter is measured from phalloidin-labelled samples. (B) Automated detection of sarcomeric repeats by MyofibrilJ (Fiji plugin), through FFT- **Fast Fourier Transform.** C(i) and (ii) Nested plots showing sarcomere length and Sarcomere diameter across analyzed samples are represented. Figure-3C(i): (n=11), 3C(ii): (n=6) n is number of myofibers.

Confocal imaging revealed the sarcomere organisation of *Drosophila* flight muscle.A single myofibril containing 6-10 sarcomeres was cropped for detailed analysis, and further magnification revealed the Z and H bands (Figure 3A).

MyofibrilJ applies an FFT to the intensity profile along the myofibril (Figure 3B), transforming spatial-domain signals into frequency-domain peaks that correspond to the regular spacing of sarcomeres. Higher-frequency peaks are assigned to brighter bands (Z band) and lower-frequency peaks to darker bands (H band) by aligning sampling points along the axis (yellow dots). This way, sarcomeres are counted by identifying Z bands and aligning them vertically, yielding a peak frequency (cycles/μm) where each cycle represents a sarcomere. Sarcomere length is finally calculated by 1/ peak frequency. (Bond et al., 2022; Katti, Bleck, et al., 2022; Schmied et al., 2024)

Quantitative analysis showed that the sarcomere length in the flight muscle averaged ∼3.4–3.8 μm. In contrast, sarcomere diameter ranges from 0.09-1.8μm across fibers (Figure 3C(i) and 3(ii)). By integrating FFT-based periodicity analysis, MyofibrilJ provides a reliable and unbiased way to assess sarcomere architecture and overcoming the inaccuracies associated with manual methods and enabling effective comparisons across varying experimental scenarios.

**Figure 4. Golgi Apparatus Morphometry and Quantitative Analysis**

Confocal images of the Golgi apparatus are represented (Figure 4A(i)). To improve visualisation of the structural detail of the Golgi, Contrast enhancement was applied (Figure 4A(ii)). Generated a threshold binary 2D mask, which helps highlight the Golgi structure for further morphometric quantification (Figure 4A(iii)).

**Figure 4.**
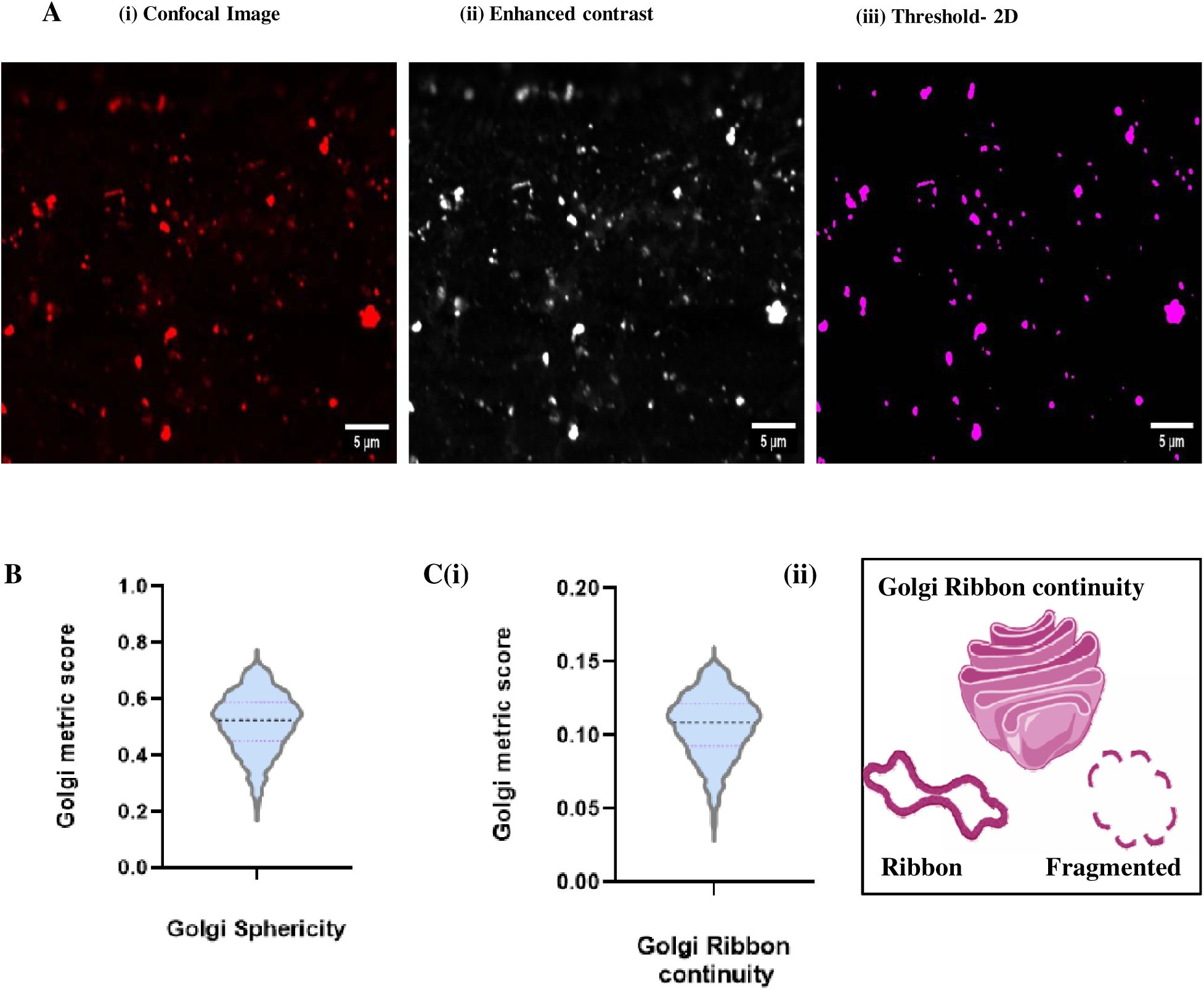
Golgi Apparatus Morphometry and quantitative analysis. A.(i) Confocal image of the Golgi apparatus. (ii) Enhanced contrast representation (iii) Thresholded mask for morphometric quantification. (B-C(i)) Violin plots representing Sphericity across Golgi data points and Ribbon continuity score Distribution of Golgi object measurements shown as median with interquartile range. Fig:4B: n=885, Fig:4C: :n=889, where n is number of golgi objects.

Quantitative analysis using Fiji/ImageJ revealed significant differences in the Golgi datasets. Ribbon continuity (RC) score describes the compactness and continuity of the Golgi ribbon; the higher the value, the more continuous the Golgi ribbon, and lower values indicate a fragmented or vesicle-like structure (Zhang & Seemann, 2024). In addition, we measured Sphericity, determining how sphere-like a structure is. In a healthy cell, they are elongated and ribbon-like like; higher sphericity indicates fragmented Golgi and lower sphericity indicates intact Golgi (Wijaya & Xu, 2024). Sphericity values suggested altered morphology consistent with fragmentation (Figure 4C(i), (ii) and 4B). Taken together, these findings indicate Golgi fragmentation, impaired ribbon continuity, and abnormal organelle morphology.

**Figure 5. Differential distribution and Volume of ER in Drosophila leg and jump muscles.**

To investigate quantitative differences in ER organization between muscle types, we compared *Drosophila* leg and jump muscles using images acquired with a confocal microscope. Perinuclear ER and regularly patterned ER structures extending along the myofibrillar axis were observed (Figure 5A, B). In leg muscles, ER distribution was limited to small punctate structures surrounding nuclei (Figure 5A), whereas jump muscles exhibited a strikingly expanded myofibrillar ER network (Figure 5C).

**Figure 5.**
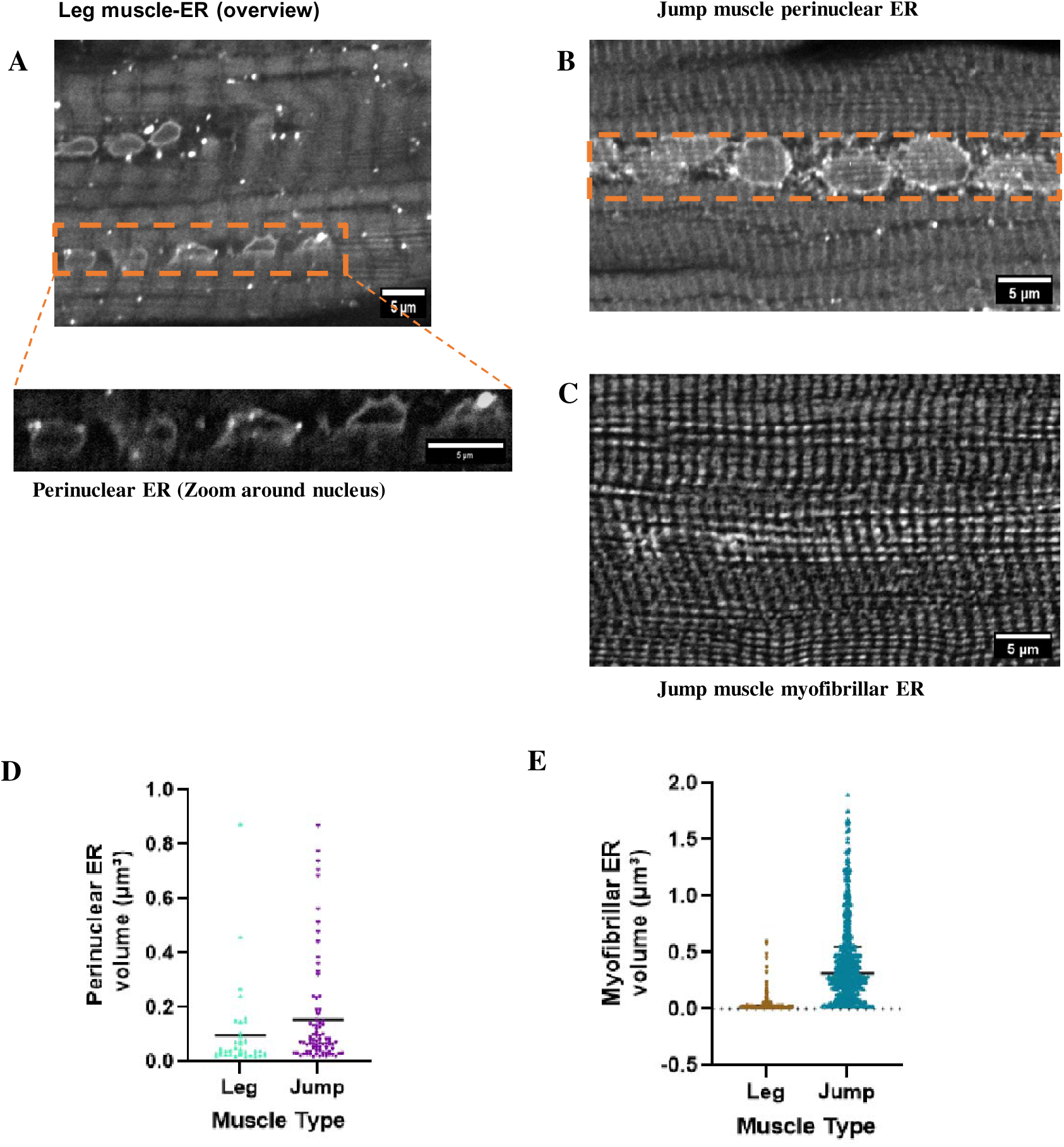
Quantification, differential distribution and Volume of ER in Drosophila leg and jump muscles. (A) Confocal micrograph showing ER organisation in leg muscle fibres. Enlarged view highlights small perinuclear ER structures (top right). (B) Perinuclear ER depiction in jump muscles(C)Dense myofibrillar ER networks are aligned in the jump muscle. (D) Quantification of Perinuclear ER volume shows significantly higher enrichment in jump muscles compared to leg muscles. (E) Quantification of myofibrillar ER shows the distribution of ER number and volume in jump and leg muscles. Data are presented as mean. Group A (n=39), Group B (n=81), Figure-5E: Group A (n=190;mean=0.04950), Group B (n=664;mean=0.4284), (n) indicate the total number of ER structures analyzed

Quantitative image analysis in Fiji revealed a significant increase in ER volume in jump muscles compared with leg muscles. Perinuclear ER volume was moderately elevated in jump muscles, whereas myofibrillar ER volume showed a highly significant expansion in jump muscles compared to leg muscles (****p < 0.0001; Figure 5E). These findings indicate that ER architecture is specialized to muscle type. The enriched perinuclear ER in jump muscles likely reflect enhanced roles in protein synthesis and calcium handling. At the same time, the extensive myofibrillar ER network may facilitate efficient excitation-contraction coupling and rapid Ca² dynamics, consistent with the mechanical demands of jump muscles (Sun et al., 2024).

**Figure 6. Imaging modality influences quantitative analysis of the cellular organelle network organization**

To evaluate how imaging modality influences quantification of subcellular structures, we compared mitochondrial and actin morphology across confocal, widefield, and super-resolution microscopy. We noticed that both the number of fragmented mitochondria and the number of actin branches were much higher in the super-resolution images than in confocal and widefield (Figure 6A).

**Figure 6.**
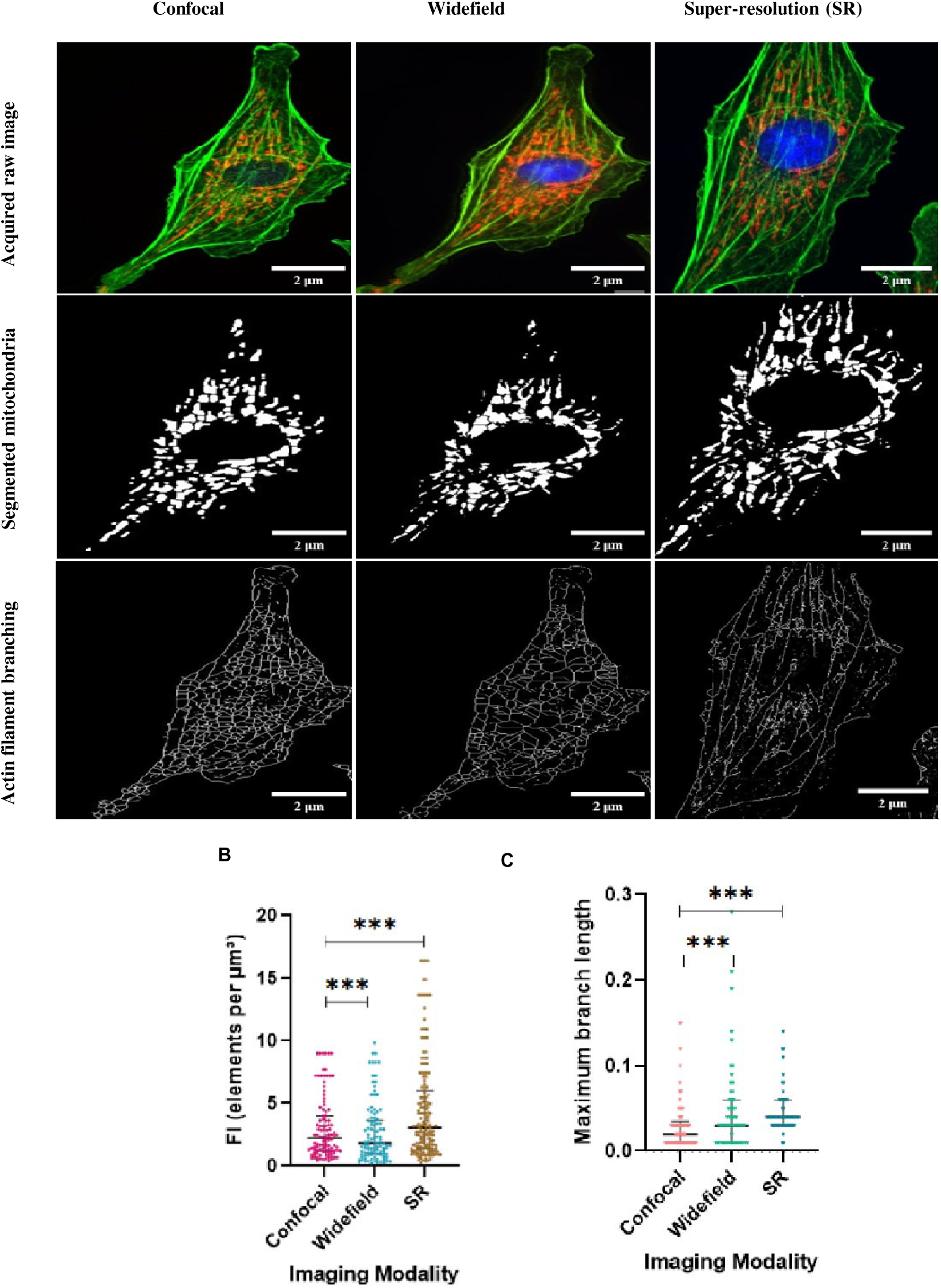
Imaging modality influences quantitative analysis of the cellular organelle network organisation. (A)Representative images of cells labelled for mitochondria (red-RFP), actin cytoskeleton(green-GFP), and nuclei(blue-DAPI), acquired using confocal(top), widefield(middle), and Super-resolution microscopy (SR)(bottom). Binary mask of mitochondria (middle) and skeletonized network representations(right) were generated in Fiji for each modality, enabling fragmentation and branching analysis. (B) Quantification of fragmentation index (FI, elements per µm³) revealed significantly higher detection of mitochondria fragments in SR compared to confocal and widefield microscopy (***p < 0.001). (C) Measurement of maximum branch length showed significantly greater detection in SR compared to confocal and widefield. Data are presented as medians with interquartile ranges, Statistical comparisons among groups were performed using Kruskal–Wallis test, n indicates the number of analyzed objects per group (Fig. 6B: Group A, n = 108; Group B, n = 106; Group C, n = 164; Fig. 6C: Group A, n = 85; Group B, n = 53; Group C, n = 51).

Quantitative analysis using Fiji revealed significant differences between modalities. The fragmentation index (FI) was significantly higher when quantified from SR images than from confocal or widefield images (Figure 6B), reflecting SR’s improved ability of to resolve small mitochondrial elements. Similarly, maximum branch length was significantly greater in SR (Figure 6C), suggesting enhanced detection of fine actin filaments extension. This doesn’t necessarily mean there are more mitochondria or branches; it’s more about how clearly one can analyze them. SR imaging gives us a much sharper view; thin filaments that might be missed or blurred together in lower resolution images become visible and countable. In contrast, confocal and especially widefield images can make small structures appear fused or even invisible, leading to underestimation during analysis (Hardo et al., 2024). Overall, this highlights how much our choice of imaging technique affects what we can measure – and it reminds us to interpret data like fragment counts carefully when comparing across different platforms.

**Figure 7. Workflow illustrating image preprocessing and analysis in Fiji.**

**Figure 7.**
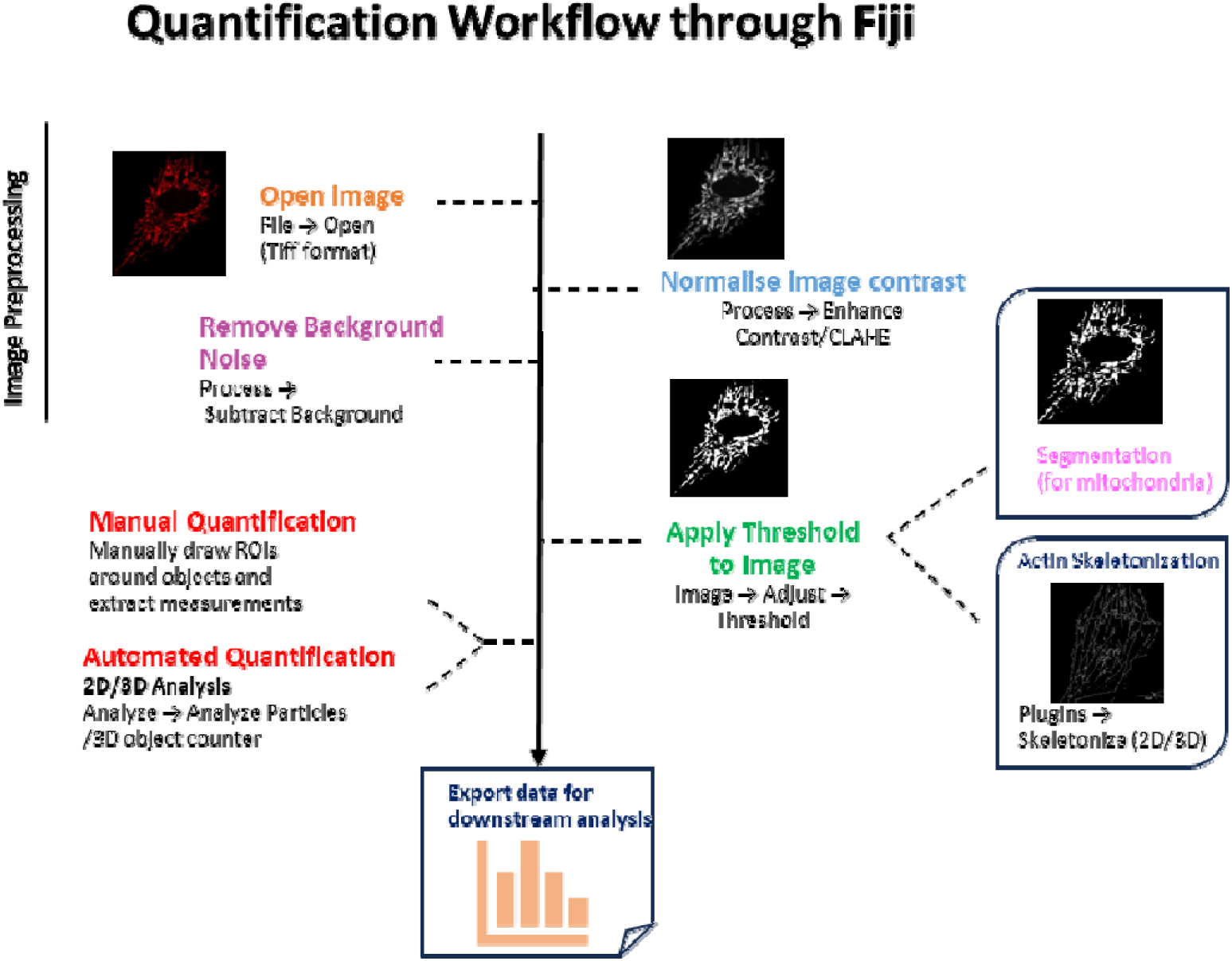
Workflow illustrating image preprocessing and analysis in Fiji.

First, raw multichannel confocal images in TIFF format are imported into Fiji. Image Preprocessing improves signal quality and ensures reliable quantification. The Subtract Background function reduces background noise from non-specific fluorescence and imaging artifacts. This separation enhances the true biological signal. To improve feature visibility and control intensity variations across samples, local contrast normalization is applied with the enhanced contrast or Contrast Limited Adaptive Histogram Equalization(CLAHE) function (J et al., n.d; Bourque et al., 2022)

After preprocessing, images undergo thresholding to covert to a binary representation. Threshold values are consistently adjusted across experimental conditions for unbiased segmentation. This process helps identify discrete organelle structures. For Example, mitochondrial analysis thresholded images create segmented mitochondrial masks. These marks are then used to extract morphometric parameters, such as shape and object count. Meanwhile, the actin cytoskeleton structure is analyzed using skeletonization algorithms (2D or 3D), which allow the quantification of filament length, branching, and network organization.

Quantification can be carried out using both manual and automated methods, depending on the analysis needs. Manual quantification requires drawing regions of interest (ROIs) around chosen structures to obtain specific measurements. Automated quantification uses particle analysis and 2D/3D object counting for high-throughput and consistent analysis across large datasets. All measurements are exported from Fiji for further statistical analysis and visualiztion, ensuring standardized and reproducible quantification of organelle morphology and spatial organiztion (Schindelin et al., 2012)

Quantitative parameters extracted to describe organelle morphology and organization. All measurements are derived from the pixel-level information and converted to physical information.

1. Area(A) - Area measurements are calculated by counting all the pixels inside a segmented object. The result is given in calibrated units (µm²).
2. Perimeter(P) - Refers to the total length of the boundary of a segmented object(µm).
3. Volume(V) - The total voxel count per object multiplied by calibrated voxel dimensions, enabling accurate volumetric analysis (Δx × Δy × Δz).
4. Circularity(C) - A Measure that shows how closely an object resembles a perfect circle. It is calculated using the object’s area and perimeter: 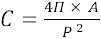
5. Aspect ratio (AR) - Ratio of the major axis length to the minor axis length of the object.
6. Form Factor (FF) - A measure of shape regularity and is calculated as: 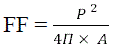
7. Inter-mitochondrial distance (IMD)- Mean Euclidean distance between centroids of neighboring mitochondria. It measures mitochondrial organization and clustering within the cytoplasm. (Courtenay et al., 2020;Neikirk, Lopez, et al., 2023b)

## Discussion

EM studies have shown how crucial it is to accurately measure organelles. This protocol advances the field by placing these ideas within a light microscopy-based framework. This workflow and protocol are applicable to cellular organelles, whereas earlier work mainly focused on ultrastructural datasets(Neikirk et al., 2023). This protocol describes a unified workflow for confocal, widefield, and super-resolution analysis, enabling high-precision quantification. Additionally, by offering validated pipelines for mitochondria, ER, Golgi, and myofibrils, and depicting how imaging methods affect quantitative results, this protocol introduces new methods and concepts that were not previously covered. Overall, these elements support extending previous frameworks by providing a standardized, accessible way to quantify organelles that can be easily adopted across different biological systems.

## Recent Methodological Advances and Future Directions

### AI-Powered Analysis Revolution

The integration of artificial intelligence has fundamentally transformed the capabilities for quantifying organelle. Deep learning approaches now achieve human-level accuracy for complex segmentation tasks while processing datasets orders of magnitude faster than manual analysis. The COSEM project demonstrated automated segmentation of 35 organelle classes, creating comprehensive cellular atlases, which were previously impossible with traditional methods (Heinrich et al., 2021; Xu et al., 2021). Weakly supervised learning addresses the fundamental challenge that biological discovery often lacks ground truth data. New paradigms enable AI-powered analysis of previously uncharacterized cellular structures, accelerating scientific discovery (Carsten et al., 2025).

### Super-Resolution Integration

Expansion microscopy has democratized super-resolution capabilities, enabling 30 nm effective resolution on standard confocal microscopes. This breakthrough makes nanoscale organelle imaging accessible to researchers worldwide without specialized equipment investments. MINFLUX approaches push resolution limits to 2-5 nm, enabling molecular-scale analysis within organelles (Carsten et al., 2025). These techniques bridge the gap between light microscopy and structural biology approaches (Moosmayer et al., 2024; Scheiderer et al., 2025; Schleske et al., 2024).

### Multi-Modal Integration

Correlative light-electron microscopy workflows now provide automated registration between imaging modalities, validating light microscopy quantification against ultrastructural gold standards. Advanced protocols maintain fluorescence throughout EM processing, enabling direct correlation of dynamic and structural measurements. Mass spectrometry integration adds molecular identification capabilities to spatial imaging data. Ramanomics platforms enable chemical fingerprinting of individual organelles, providing comprehensive functional characterization. The convergence of these technological advances positions light microscopy-based organelle quantification as a cornerstone methodology for 21st-century cell biology. Researchers can now address fundamental questions about cellular organization with unprecedented precision, throughput, and biological relevance, opening new frontiers in our understanding of how organelles shape cellular function in health and disease.

### Time Considerations

Complete protocol implementation typically requires 2-4 weeks, including validation. Sample preparation: 1-3 days. Image acquisition: 1-5 days, depending on throughput requirements. Analysis: 2-7 days for comprehensive quantification. Critical Success Factors: Rigorous experimental design, appropriate controls, validated analysis workflows, and comprehensive statistical analysis ensure reliable, reproducible results suitable for publication in high-impact venues.

## Acknowledgements

We acknowledge all contributing authors and laboratory members for their valuable scientific discussions, critical insights, and constructive feedback throughout the manuscript development process. Software: Schematic illustrations and conceptual diagrams were created using BioRender software (BioRender.com). We thank Mr. Ganesh Kadasoor of Evident Scientific India Ltd., Bangalore, India, for his technical assistance with the imaging.

## Author Contributions

Conceptualization S.T., H.K, H.P., A.H.J., P.K.,; Methodology S.T., H.K., H.P., A.M., P.K.; RM., Writing—Original Draft Preparation,. S.T., H.K, H.P; Writing—Review & Editing, H.K., S.T., P.K., H.P., B.G., A.M., P.V., A.H.J.,; Supervision, B.G., A.H.J., P.K..

All authors have read and agreed to the published version of the manuscript.

